# Targeting Rab-RILPL Interactions as a Strategy to Downregulate Pathogenic LRRK2 in Parkinson’s Disease

**DOI:** 10.1101/2023.11.06.565863

**Authors:** Krista K. Alexander, Yahaira Naaldijk, Rachel Fasiczka, Besma Brahmia, Tiancheng Chen, Sabine Hilfiker, Eileen J. Kennedy

## Abstract

Familial Parkinson’s disease (PD) is frequently linked to multiple disease-causing mutations within Leucine-Rich Repeat Protein Kinase 2 (LRRK2), leading to aberrant kinase activity. Multiple pathogenic effects of enhanced LRRK2 activity have been identified including loss of cilia and centrosomal cohesion defects. When phosphorylated by LRRK2, Rab8a and Rab10 bind to phospho-specific RILPL effector proteins. RILPL-mediated accumulation of pRabs proximal to the mother centriole is critical for initiating deficits in ciliogenesis and centrosome cohesion mediated by LRRK2. We hypothesized that Rab-derived phospho-mimics may serve to block phosphorylated Rab proteins from docking with RILPL in the context of hyperactive LRRK2 mutants. This would serve as an alternative strategy to downregulate pathogenic signaling mediated by LRRK2, rather than targeting LRRK2 kinase activity itself. To test this theory, we designed a series of constrained peptides mimicking phosphorylated Switch II derived from Rab8. These RILPL interacting peptides, termed RIP, were further shown to permeate cells. Further, several peptides were found to bind RILPL2 and restore ciliogenesis and centrosomal cohesion defects in cells expressing PD-associated mutant LRRK2. This research demonstrates the utility of constrained peptides as downstream inhibitors to target pathogenic LRRK2 activity and may provide an alternative approach to target specific pathways activated by LRRK2.

## 1. INTRODUCTION

Parkinson’s disease (PD) is a neurodegenerative disease that may be initiated sporadically or through genetic changes [1]. While the majority of PD cases are sporadic, approximately 10% are familial with *LRRK2* being the most frequently mutated gene [2]. Further, *LRRK2* mutations have also been linked to sporadic PD [1]. The gene product, Leucine-Rich Repeat Kinase 2 (LRRK2), is a large, multidomain protein. The preponderance of mutations linked to PD are localized to the enzymatic domains of the protein, including the kinase domain and the GTPase-active Roc domain [3]. Further, these mutations were all found to increase the kinase activity of LRRK2 [4, 5]. However, the underlying mechanisms of LRRK2 activation and regulation remain poorly understood. Significant efforts have been put forth to develop ATP-competitive kinase inhibitors that are selective for LRRK2 [6-10], however, long-term inhibition of LRRK2 kinase activity by many of these catalytic inhibitors leads to altered LRRK2 localization and major side effects including morphological alterations in lung and kidney [11, 12]. Due to these effects, no LRRK2 inhibitors have yet received FDA approval.

When LRRK2 kinase activity is upregulated due to disease-associated mutations, the phosphorylation of several Rab GTPases is also upregulated [4, 13]. Humans express over 60 Rab proteins that regulate intracellular membrane trafficking [14, 15]. Rab proteins are members of the Ras superfamily of small GTPases that switch between the active GTP-bound state and the inactive GDP-bound state, which involves conformational changes of two regions called switch I and switch II that are modulated by various regulatory proteins. In their GTP-bound active state, Rab proteins bind to their effector proteins by interacting with their switch II regions [16]. Several Rab proteins have been found to be physiological substrates of LRRK2, including Rab8a and Rab10, where LRRK2 phosphorylates a conserved Ser or Thr residue in the switch II region [4].

After phosphorylation by LRRK2, a pool of Rab8a and Rab10 were found to concentrate around the centrosomes and bind to a class of phospho-specific effector proteins, termed Rab Interacting Lysosomal Protein-Like (RILPL) proteins [17]. Higher vertebrates have two homologous forms of RILPL: RILPL1 and RILPL2 [18]. The interaction between phosphorylated Rab8a and Rab10 and RILPL proteins at the mother centriole/ciliary base was found to cause both ciliary and centrosomal cohesion defects in various cell types *in vitro* [13, 17, 19-22]. Physiologically, these pRab-RILPL complexes also cause deficits in centrosome cohesion including in primary or immortalized lymphocytes from LRRK2-PD patients as compared to healthy controls [20, 23, 24]. Further, these effects can be reversed using ATP-competitive LRRK2 inhibitors, demonstrating the instrumental role of hyperactive LRRK2 in initiating ciliogenesis and cohesion defects. Cilia are considered fundamental regulators for a variety of cell processes, and ciliary deficits have also been observed in the brain of mutant LRRK2-KI (knock-in) mice where they may interfere with a neuroprotective circuit aimed at maintaining dopaminergic cell health [17, 25].

At the molecular level, a crystal structure of the phosphorylated form of Rab8a was previously solved in complex with the RILP homology domain (RH2) of RILPL2, exposing the protein-protein interface mediating this interaction (**Figure 1A**) [26]. This structure reveals how the phosphorylated form of switch II from Rab8a binds to RILPL2 and highlights several critical contact points in the interface including RILPL2 residues R130, R132, F133, E137 and K149 (**Figure 1B**) [26]. These residues are conserved in other homologous scaffolding proteins, including RILPL1 and JIP4, which were previously found to bind phosphorylated Rab10 [17, 26].

**Figure 1.**
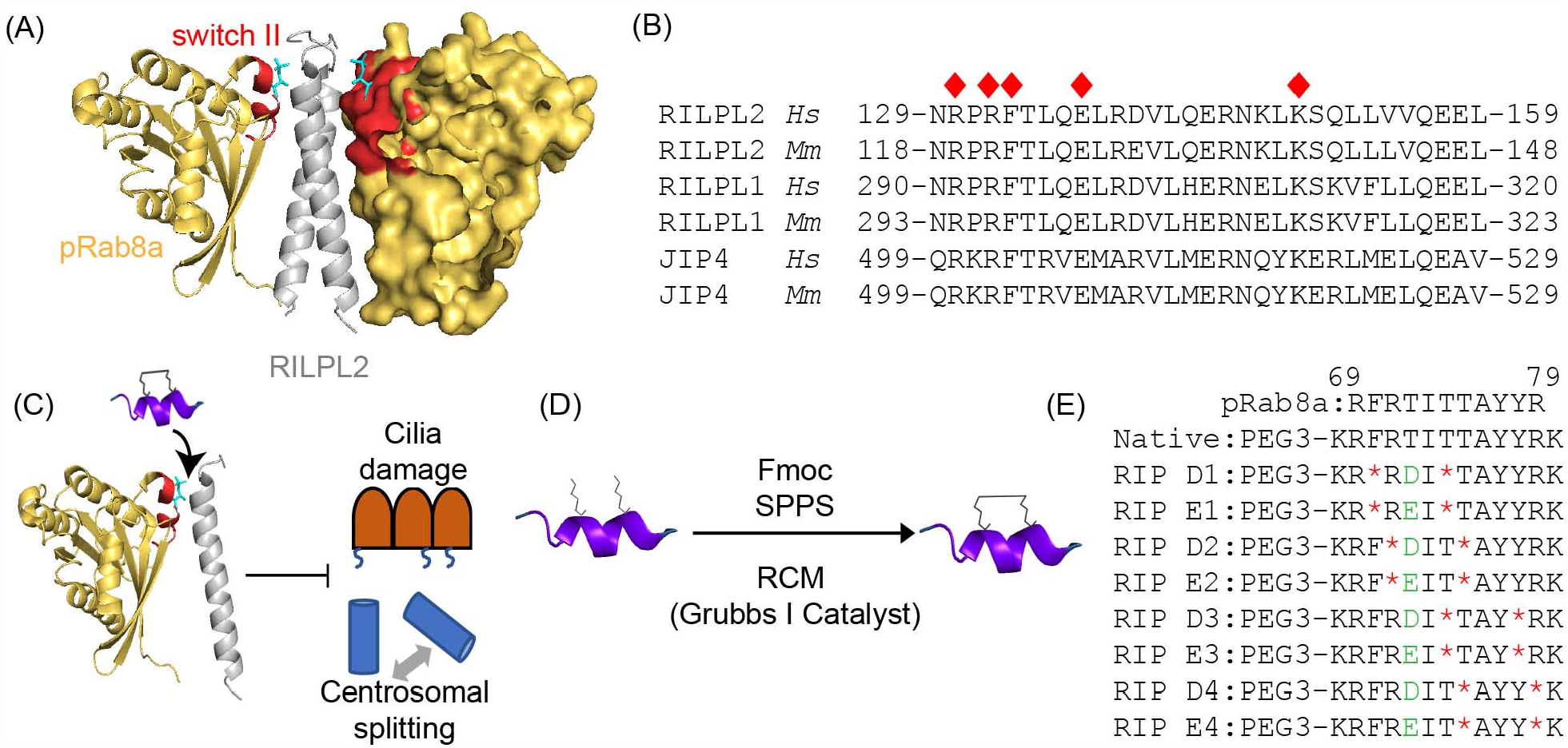
Design of a constrained peptide library targeting the Rab8a:RILPL2 interface. (A) Crystal structure showing the interactions between RILPL2 (grey) and pRab8a (yellow). Switch II of Rab8a is highlighted in red with the phosphate group shown in teal. (B) Sequence alignment of human and murine versions of effector proteins RILPL2, RILPL1, and JIP4. Key residues in the Rab8a:RILPL2 interface are marked with red diamonds. (C) Cartoon illustration demonstrating how constrained peptides mimicking switch II may disrupt this PPI, thereby downregulating downstream effects such as cilia loss and centrosomal cohesion defects. (D) Peptides were synthesized using standard Fmoc solid-phase synthesis (SPPS). The olefinic amino acids were cyclized using ring-closing metathesis (RCM) chemistry while on solid support. (E) The peptide library was designed to mimic switch II. Phosphate mimetics (either Asp or Glu) were incorporated for each staple position and are shown in green. Red asterisks represent the positioning of the non-natural amino acid (S)-N-Fmoc-2-(4-pentenyl) alanine.

Prior work demonstrated that the Rab/RILPL protein-protein interface (PPI) could perhaps serve as a target to ameliorate the downstream effects of aberrant LRRK2 activity. In this study, a C-terminal fragment of RILPL1 was expressed in cells that could bind to phosphorylated Rab proteins but was unable to localize to the centrosome. This fragment reverted the cohesion defects mediated by pathogenic LRRK2 by redistributing phosphorylated Rab10 proteins away from the centrosome without altering the total levels of phosphorylated Rab proteins [21]. These critical proof-of-principle experiments indicate that targeting the pRab-RILPL interaction may serve as a feasible approach to revert downstream cellular defects due to pathogenic LRRK2 without altering LRRK2 kinase activity.

As a strategy to modulate LRRK2 pathogenesis, we sought to target the PPI between Rab8-RILPL2 (**Figure 1C**). To accomplish this, we developed a series of all-hydrocarbon-constrained peptides mimicking switch II derived from Rab8a (**Figure 1D, E**). These constrained peptides were found to be helical and cell-permeant. In addition, several peptides were found to bind their intended target, RILPL2. Further, the peptides were found to reverse ciliogenesis and centrosomal cohesion defects in cells in the presence of a pathogenic mutant form of LRRK2, R1441C, to levels comparable to the small molecule LRRK2 inhibitor, MLi-2. Although no inhibitors currently exist that target downstream pathogenic effects mediated by LRRK2 signaling, this work highlights the utility of constrained peptides to inhibit the pathogenic effects of LRRK2 and may serve as an alternative strategy to target specific pathways activated by LRRK2.

## 2. MATERIALS AND METHODS

### 2.1 Peptide Synthesis

N-*α*-Fmoc amino acids and rink amide MBHA resin used in this work were purchased from Novabiochem. (S)-N-Fmoc-2-(4-pentenyl) alanine (S_5_), Grubb’s I catalyst, and FAM (5(6)-carboxyfluorescein) were purchased from Sigma-Aldrich. D-biotin was purchased from GoldBio. All solvents and other liquid reagents were purchased from Fisher Scientific: anhydrous diethyl ether, piperidine, N-methyl pyrrolidinone (NMP), 1,2-dichloroethane (DCE), dimethylformamide (DMF), dimethyl sulfoxide (DMSO), N, N-diisopropyl ethylamine (DIEA), trifluoroacetic acid (TFA), triisopropylsilane (TIS), methyl-tert-butyl ether (MTBE), dichloromethane (DCM). O-(1H-6-Chlorobenzotriazole-1-yl)-1,1,3,3-tetramethyluronium hexafluorophosphate (HCTU) was purchased from Oakwood-Chemical. Methanol was purchased from Honeywell.

Peptides were synthesized on rink amide MBHA resin (Novabiochem). Synthesis was carried out using standard solid-phase peptide synthesis (SPPS) where the Fmoc protecting group was removed prior to the coupling of each amino acid (deprotecting solution: 25% piperidine in NMP). For each coupling, 10 equivalents of amino acid, 9.9 equivalents of HCTU and 20 equivalents of DIEA were used. Each deprotection step occurred for 25 minutes and each coupling step occurred for 45 minutes, with three NMP washes after each step. For S_5_ coupling steps, 4 equivalents of the olefinic amino acid were added along with 3.9 equivalents of HCTU and 20 equivalents of DIEA for 45 min. The ring-closing metathesis reaction (RCM) was performed using 0.4 equivalents of Grubb’s I catalyst dissolved in DCE for 1 h and was repeated for an additional cycle. N-terminal FAM labeling was performed by adding 2 equivalents of 5(6)-FAM dissolved in DMF, 1.8 equivalents of HCTU, and 4.6 equivalents of DIEA overnight. N-terminal biotin labeling was performed by adding 10 equivalents of biotin dissolved in DMF and DMSO at a 1:1 ratio, 9.9 equivalents of HCTU, and 20 equivalents of DIEA. Peptides were cleaved from resin using 95% TFA (v/v), 2.5% (v/v) TIS, and 2.5% (v/v) water. This solution was allowed to rotate for 4-5 h, followed by filtration into ice-cold MTBE. Peptides were then centrifuged at 2800 rpm for 30 minutes, the supernatants were removed, and the pellet was air-dried overnight. Prior to characterization and purification, peptides were redissolved in 1 mL methanol and filtered through a 45 μm syringe.

Peptide Purification and Quantification

Peptides were analyzed by LC/MS using electrospray-ionization mass spectrometry (ESI-MS, Agilent 6120 Single Quadrupole). Peptides were purified using reversed-phase high-performance liquid chromatography (RP-HPLC, Agilent 1250). using a semi-prep Zorbax SB-C18 column.

Initial mass spectras were performed using an analytical SB-C18 column with a flow rate of 0.5 mL/min using a 10-100% gradient of acetonitrile and 0.1% TFA in water. Using the same mobile phase, purification was conducted using a semi-prep SB-C18 column with a flow rate of 4 mL/min. Final mass spectra confirmed the products for each synthesis **(Figure S1-S5)**. Peptides were quantified using either their FAM or biotin labels. For FAM-labeled peptides, absorbance was measured at 495 nm, using 10 mM Tris buffer, pH 8.0 and an extinction coefficient of 69,000 M^-1^cm^-1^. For biotin-labeled peptides, 2-hydroxyazobenzen-4’-caryboxylic acid (HABA)-avidin complex was used (Thermo Fisher). Quantification was measured at 500 nm, where the quantification of the peptide was determined by the decrease in the absorbance of the HABA-avidin complex in the presence of the peptide.

*The sequence names, amino acid residues, and molecular weight details of each peptide discussed are shown (*^*^ = S-2-(4’-pentenyl) alanine):

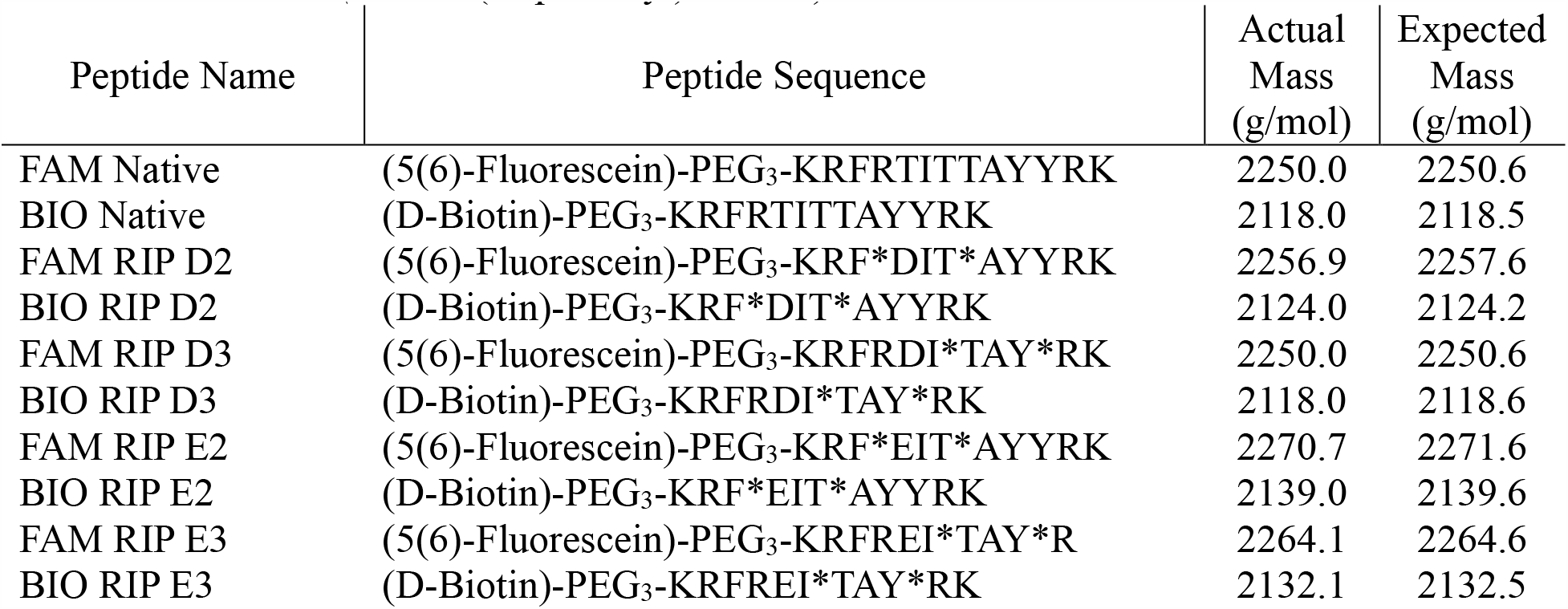

### 2.2 Cell Culture

A549 cells (ATCC, CCL185) were cultured in RPMI 1640 media supplemented with 10% fetal bovine serum (FBS) and 1% penicillin-streptomycin-glutamine. The RPMI media was purchased from Corning, the FBS from HyClone, and the penicillin from Gibco. Phosphate-buffered saline (PBS) was purchased from Lonza.

### 2.3 Cell Uptake Experiments

A549 cells were plated at 20,000 cells per well and grown in supplemented RPMI media on CC^2^-treated chamber plates for 48 hours prior to peptide treatments. Cells were then treated with 2.5 μM of individual FAM-labeled peptides for 8 hours. After treatment, cells were washed three times in PBS then permeabilized and fixed in a solution containing 4% paraformaldehyde and 0.1% Triton X-100 in PBS. Cells were washed three additional times with PBS followed by nuclear staining with 0.1 μg/mL Hoechst 34580 stain for 5 minutes. Cells were then mounted with PermaFluor (PerkinElmer) prior to imaging. Cell uptake was imaged on a Zeiss fluorescent microscope (Olympus IX71) at 40X magnification.

### 2.4 Pull-down Experiments

A549 cells were cultured and grown in 10 cm dishes at 2 million cell density to achieve full confluency. Cells were then washed once in ice-cold PBS and lysed using trypsin prior to being centrifuged. The supernatant was removed, and cell pellets were kept on ice and resuspended in a non-denaturing lysis buffer (20 mM Tris HCl pH 8, 137 mM NaCl, 10% glycerol, 1% nonidet P-40 (NP-40), 2 mM EDTA). This mixture was rotated at 4 °C for 30 minutes, then centrifuged for 10 minutes at 10,000 rpm. The supernatant was collected, and biotin-labeled peptides were added at a final concentration of 2.5 μM to the cell lysates. This cocktail was allowed to rotate at 4 °C for 4 hours before adding 50 μL of streptavidin agarose resin (Millipore) overnight with continued rotation at 4 °C. The next day, cells were centrifuged, and the pellet was washed twice in tris buffer before being resuspended in 50 μL 2X Laemmli sample buffer. The immunoblotting conditions previously described were again applied here to generate the blots. A 1:500 dilution was applied for all primary antibodies (RILPL2, GeneTex (GTX121000); RILPL1, Abcam (ab302492); JIP4, Cell Signaling (5519)) and 1:10,000 for goat-Anti-Rabbit 800CW/680RD secondary antibody (Li-COR). Blots were then imaged on the Li-COR imager. Each treatment was independently performed in triplicates. Western blots were quantified as a ratio of average input to treatment. Image Studio software was used to quantify the pulldown binding. For each treatment (RILPL2, RILPL1, or JIP4), the exposures were consistent to quantify and normalize input to treatment expression. Statistical analysis was performed using GraphPad Prism. One-way ANOVA and Dunnett’s multiple comparisons test within the Prism software were used to generate p-values based on standard deviation across the triplicates for each treatment. The statistical significance of each treatment was generated compared to the DMSO treatment. For each statistical analysis, the following abbreviations were used: n.s. = not significant, *p < 0.05, ^**^p < 0.01, ^***^p < 0.001, and ^****^p < 0.0001.

### 2.5 Circular Dichroism

Peptides were diluted in water and were analyzed using a 1.00 mm QS cuvette (Starna Cells). A JASCO J-710 circular dichroism spectrometer and JASCO Spectra Manager software were used to analyze the results using the following parameters: 100 mdeg sensitivity and a scanning mode of 50 nm/min. After collecting each peptide sample, the blank sample was subtracted from each spectra and smoothed.

### 2.6 Proteolytic Stability

CD1 mouse serum was purchased from Innovative Research Inc. Benzyl alcohol was purchased from Fisher. For each peptide treatment, a cocktail containing 0.2 mM of peptide, 50% (v/v) mouse serum, 0.4% (v/v) benzyl alcohol, and 15% (v/v) DMSO in PBS was used. Each cocktail was incubated at 37 °C with agitation for 6 h. At time intervals of 0, 2, 4 and 6 hours, an aliquot of the solution was collected and quenched in an equal volume of 0.1% TFA in acetonitrile. This solution was then centrifuged at 14,000 rpm for 5 min and the supernatant was collected for ESI-MS analysis. The same reversed-phase gradient previously described for mass spectra analysis was applied. Experiments were performed in triplicate for each time point. The percentage of peptide remaining for each time point was measured as a ratio of peptide relative to the internal standard (benzyl alcohol). The values were normalized to time 0 h for each peptide. Statistical analysis was performed using GraphPad Prism software to calculate the standard deviation.

### 2.7 Centrosomal Cohesion Determination

Homozygous LRRK2-R1441C knock-in murine embryonic fibroblasts (R1441C-LRRK2 MEFs) were generated from mice at E12.5 and spontaneously immortalized by prolonged passaging, and they were a generous gift from Dario Alessi [13]. R1441C-LRRK2 MEFs were chosen as they express high endogenous levels of LRRK2 and display centrosome cohesion and ciliogenesis defects as compared to littermate wildtype MEFs which are reverted by the LRRK2 kinase inhibitor MLi2 [21, 27, 28]. Cells were grown in full medium consisting of DMEM containing high glucose (Gibco, 11960-044), 10% fetal bovine serum (Gibco, 10438-026), 1 mM sodium pyruvate (Gibco, 11360-070), non-essential amino acids (Gibco, 11140-050), 2 mM L-glutamine (Gibco, 25030-081), 100 U/ml penicillin and 100 μg/ml streptomycin (Gibco, 15140-122). Cells were passaged at around 80-90% confluency to a ratio of 1:10 for general maintenance, with media exchanged every other day. Cells were grown at 37 ºC and 5% CO_2_ in a humidified atmosphere and were regularly tested for mycoplasma contamination.

To determine centrosome cohesion, cells were seeded onto coverslips in full medium, and at around 80% confluency were treated with 200 nM MLi2 (Abcam, ab254528), with the equivalent volume of DMSO (< 0.1% v/v), or with the indicated concentrations of FAM-labeled peptides for 4 h, followed by a rinse with full medium. Cells were fixed with 4% paraformaldehyde (PFA) in PBS for 20 min at RT, followed by ice-cold MeOH for 10 min at -20 ºC. Cells were permeabilized in 0.5% Triton-X100/PBS (Tx-PBS) for 10 min and blocked in 0.5% BSA in 0.5% Tx-PBS for 1 h at room temperature before incubation with mouse monoclonal anti-γ-tubulin antibody (Abcam, ab11316, 1:1000) in 0.5% BSA in 0.5% Tx-PBS overnight at 4 ºC. The next day, coverslips were washed three times for 10 min with 0.5% Tx-PBS, followed by incubation with secondary antibody (Alexa647-conjugated anti-mouse, Invitrogen, 1:1000) in 0.5% Tx-PBS for 1 h at room temperature. Coverslips were washed three times in 0.5% Tx-PBS, rinsed with PBS, dried at RT, and mounted in mounting media with DAPI (Vector Laboratories).

Images were acquired on an Olympus FV1000 Fluoview confocal microscope using a 60x 1.2 NA water objective lens. Images were collected using single excitation for each wavelength separately. Around 10-15 optical sections of randomly selected areas were acquired with a step size of 0.5 μm, and maximum intensity projections of z-stack images analyzed and processed using ImageJ. As previously described [27, 29], the distance between two centrosomes was measured for around 100 cells per condition and experiment, and centrosomes were scored as being split when the distance between them was > 5 μm. Mitotic cells were excluded from the analysis in all cases.

### 2.8 Ciliogenesis determination

Cells at around 80-90% confluency were serum-starved for 24 h in either the presence or absence of DMSO, 200 nM MLi2 or the indicated concentrations of FAM-labelled peptides as indicated. Please note that the concentration range of peptides for a dose-response curve for ciliogenesis was less than that employed for a dose-response curve for cohesion, possibly due to differences in media composition and incubation times (24 h for ciliogenesis in medium without serum, 4 h for cohesion in medium with serum).

Coverslips were fixed, permeabilized and blocked as described above, and incubated with rabbit polyclonal anti-Arl13b (Proteintech, 177-11-1-AP, 1:400) and mouse monoclonal anti-γ-tubulin (Abcam, ab11316, 1:1000) primary antibodies in 0.5% BSA in 0.5% Tx-PBS overnight at 4 ºC. The next day, coverslips were washed three times for 10 min with 0.2% Tx-PBS, followed by incubation with secondary antibodies (Alexa647-conjugated anti-mouse and Alexa594-conjugated anti-rabbit, Invitrogen, 1:1000) in 0.5% Tx-PBS for 1 h at room temperature. Coverslips were washed three times in 0.5% Tx-PBS, rinsed with PBS and mounted in mounting media with DAPI (Vector Laboratories).

Images were acquired on an Olympus FV1000 Fluoview confocal microscope using a 60x 1.2 NA water objective lens as described above, and maximum intensity projections of z-stack images were analyzed and processed using ImageJ. Cilia were scored from 200 cells per condition and experiment based on an Arl13b-positive cilium emanating from a γ-tubulin-positive ciliary base [19]. For select experimental conditions, quantification of ciliogenesis was performed by an additional observer blind to conditions, with identical results obtained in both cases.

## 3. RESULTS AND DISCUSSION

### 3.1 Constrained Peptide Library Design

Based upon the crystal structure of Rab8a in complex with the RILPL2 RH2 domain (PDB: 6RIR), a library of constrained peptides was designed using the switch II region of Rab8a (**Figure 1D, E**). The S-2-(4’-pentenyl) alanine residues were introduced in *i, i*+4 positions to stabilize individual helical turns of the sequence. In addition, these olefinic amino acids were placed in positions that were anticipated to orient the hydrocarbon staple onto the solvent-exposed face of the helix. Single lysine residues were added to the N- and C-terminus to increase the overall net charge of the peptides. Further, PEG_3_ was included at the N-terminal of each sequence to promote water solubility. In addition, since the phosphorylation of Thr 72 was found to be critical for promoting interactions with RILPL2, Asp or Glu phosphomimetics were substituted at this position. Peptides were synthesized using standard Fmoc solid phase peptide synthesis and the ring-closing metathesis (RCM) reactions were performed on-resin. These peptides, termed RILPL Inhibitory Peptides (RIP peptides), were subsequently used for biochemical characterization.

### 3.2 Constrained Peptides Permeate Cells and are Helical

To determine whether the constrained peptides could permeate cells, FAM-labeled peptides were incubated with A549 cells, which were chosen since they intrinsically express high levels of LRRK2. Cells were treated with 2.5 μM of individual peptides for 8 hours prior to fixation and imaging **(Figure 2A, S6)**. In contrast to the native peptide, all constrained peptides in the library were found to permeate cells and were detected throughout the nucleus and cytoplasm. Of note, peptide RIP D3 was found to particularly localize to puncta within the cell, likely due to localization within intracellular vesicles. Four peptides, RIP D2 and E2 and RIP D3 and E3, were chosen for subsequent experiments due to their superior extent of cell permeation.

**Figure 2.**
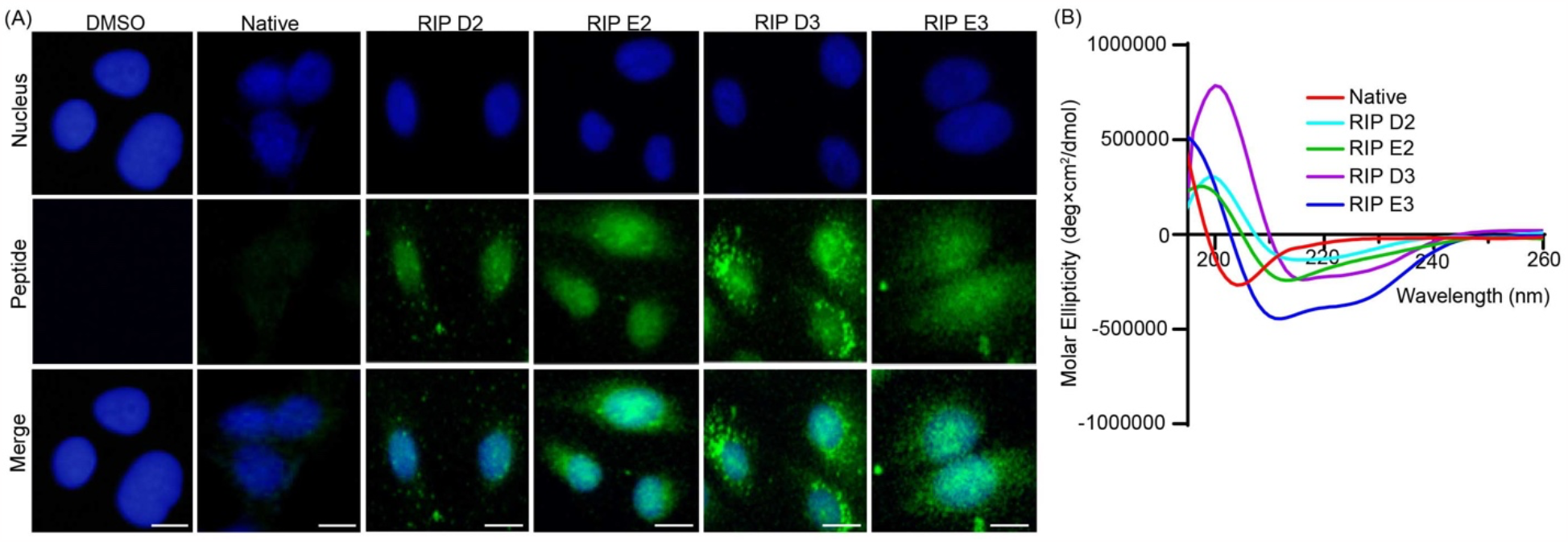
Constrained peptides permeate cells and are helical. (A) A549 cells were treated with 2.5 μM of FAM-labeled RIP peptides for 8 hours, followed by imaging. Cell images are representative of n ≥ 200. All constrained peptides were found to permeate cells and localize diffusely throughout the cytoplasm and nucleus, while the unconstrained native peptide did not. The scale bar represents 10 μm. (B) Circular dichroism analysis of FAM-labeled peptides (native, RIP D2, RIP E2, RIP D3, and RIP E3). The constrained peptides demonstrated alpha-helical content with minima at 208 and 222 nm, while the unconstrained native peptide did not. CD measurements were performed in triplicate.

Since the peptides were designed to mimic the switch II helix of Rab8a, we next wanted to assess whether this secondary structure was reinforced with the introduction of the hydrocarbon staple. The peptides were analyzed by circular dichroism (CD) **(Figure 2B)**. From this analysis, each of the stapled peptides tested was found to have minima at 208 and 222 nm, demonstrating alpha-helical content. On the other hand, the non-constrained native peptide sequence had minimal detectable alpha-helical content. Thus, the incorporation of the hydrocarbon staple appears to have helped reinforce an alpha-helical structure and cell permeation.

### 3.3 Constrained Peptides Bind to RILPL2 and are Resistant to Proteolytic Degradation

Next, we sought to identify whether the RIP peptides could bind their intended intracellular target, RILPL2. To test this, pull-down assays were performed using biotin-labeled versions of the RIP peptides or a DMSO control incubated with A549 cell lysates, followed by treatment with streptavidin beads. Interacting partners were analyzed by western blotting with an anti-RILPL2 antibody (**Figure 3A, B**). All the RIP peptides tested were found to interact with RILPL2 with statistical significance relative to the control, with RIP E3 demonstrating the greatest detected band intensity. Since the sequence of the RH2 domain in RILPL2 is nearly identical to RILPL1 and is also homologous to JIP4, we additionally determined whether the RIP peptides may bind to these other adaptor proteins. Additional pull-downs were performed and either RILPL1 or JIP4 interactions were detected by western blotting (**Figure 3C-F**). RIP E3 was found to also interact with RILPL1, but the other peptides tested did not have detectable interactions with this effector protein. In addition, none of the peptides detectably interacted with JIP4 as expected since JIP4 has a greater extent of sequence diversity as compared to RILPL2. Thus, while the peptides bind RILPL2 as expected, they may additionally have off-target effects by binding to RILPL1, and perhaps other homologous proteins, within the cellular environment.

**Figure 3.**
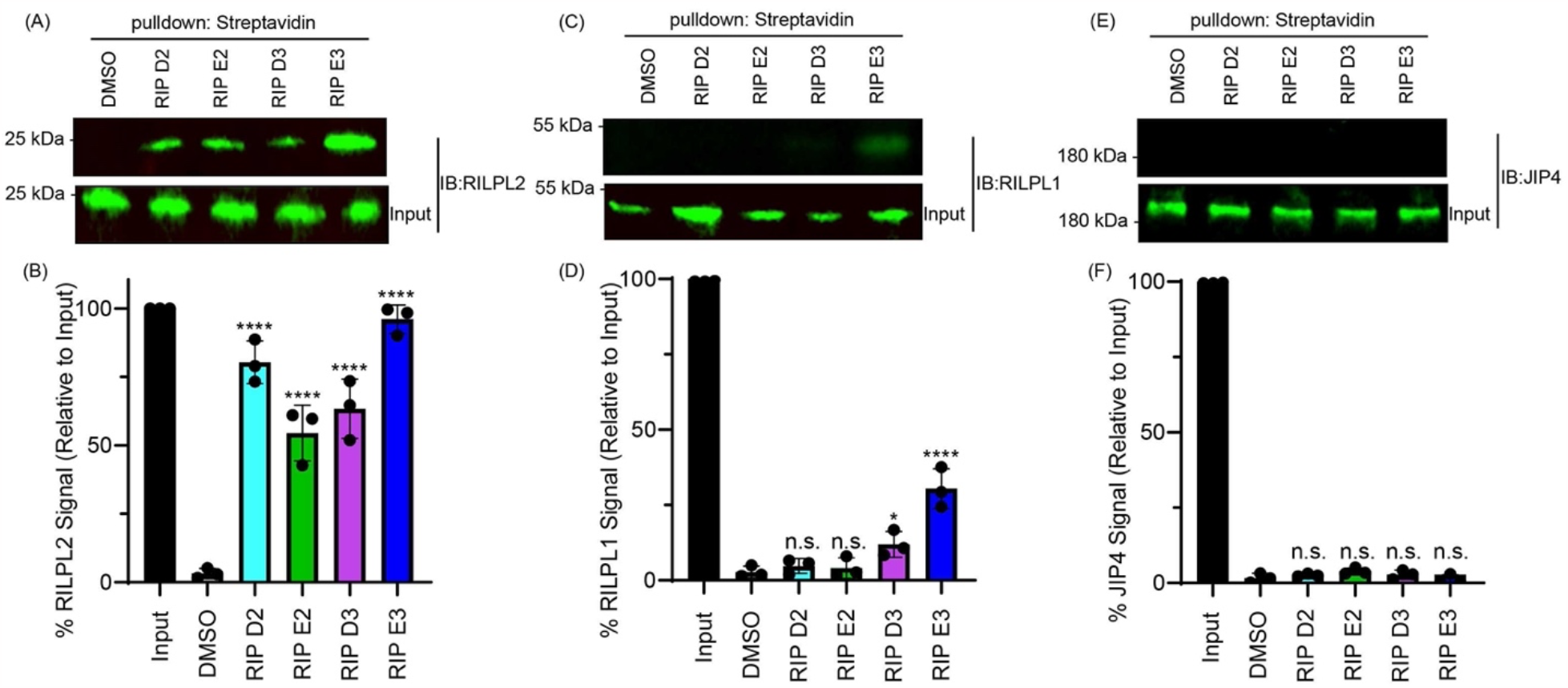
Constrained peptides bind to RILPL2. (A) Streptavidin pull-down experiments were conducted in A549 cells with 2.5 μM of individual biotin-labeled peptides. Western blotting was performed using anti-RILPL2, demonstrating that all RIP peptides interact with their intended target. (B) Quantification of RILPL2 western blots from three independent experiments. ^****^p<0.0001. (C) Pull-down experiments, followed by western blot analysis to detect RILPL1 interactions. Only RIP E3 had notable interactions with RILPL1. (D) Quantification of RILPL1 western blots from three independent experiments. *p<0.05, ^****^p<0.0001, n.s.= not significant. (E) Pull-down experiments, followed by western blot analysis to detect JIP4 interactions. None of the RIP peptides had detectable interactions with this protein. (F) Quantification of JIP4 western blots from three independent experiments. n.s.= not significant.

Further, since the RIP peptides were designed to be used in cells, proteolytic stability was measured **(Figure S7)**. To measure resistance to degradation, we incubated 0.2 mM of each peptide with mouse serum which contains a variety of proteases. This cocktail was incubated at 37 °C over a 4-hour time course. Degradation of each peptide was measured by LC/MS to analyze the loss of the parent compound over time. Benzyl alcohol was used as an internal standard for analysis. Over the time course tested, the unstapled native peptide was degraded by nearly 50%. On the other hand, all four RIP peptides tested maintained their integrity over time. Over 75% of RIP E2 was detected by 4 hours, while over 90% of RIP D2, RIP D3, and RIP E3 were still detected at this time point. These results suggest that the incorporation of the staple notably improved the proteolytic stability of the RIP peptides.

### 3.4 Peptides Restore Ciliogenesis Deficit in Cells

Disease-relevant mutant forms of LRRK2 increase kinase activity and are associated with increased phosphorylation of substrates including Rab8 and Rab10, thereby resulting in deficits in ciliogenesis. Therefore, we investigated whether the RIP peptides could reverse this effect by blocking interactions between RILPL and phosphorylated Rab proteins in knock-in MEF cells endogenously expressing a mutant form of LRRK2, R1141C, that were serum-starved for 24 hours (**Figure 4A, B**). Each RIP peptide was tested over a dosage range of 1-2.5 μM added for 24 hours. DMSO or the LRRK2 kinase inhibitor MLi-2 were used as controls. Cells were subjected to immunocytochemistry with an antibody against γ-tubulin to detect the ciliary base, an antibody against Arl13b to stain cilia and DAPI as a nuclear stain. Percentage of ciliogenesis was scored from 200 cells per condition and experiment. The greatest reversal was observed at the highest concentration tested, 2.5 μM, and this effect was nearly on par with the effects of the ATP-competitive LRRK2 kinase inhibitor, MLi-2. None of the peptides demonstrated effects on ciliogenesis at the lowest concentration tested (1 μM). This demonstrates that an allosteric approach targeting downstream signaling of mutant LRRK2 can have comparable effects to direct catalytic inhibition of LRRK2 on reverting the ciliogenesis deficit.

**Figure 4.**
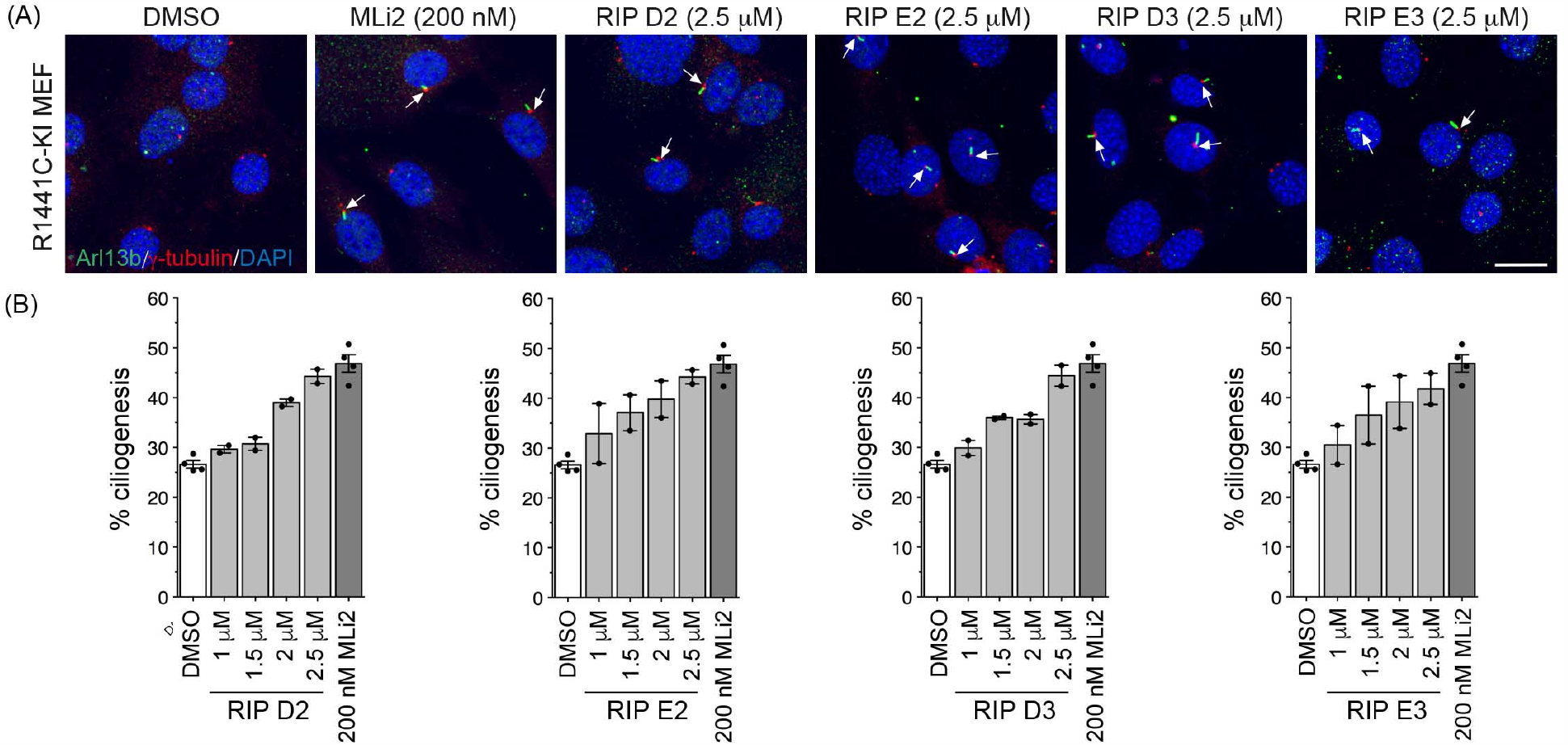
Constrained peptides revert ciliary deficits in a cell line harboring a PD-associated LRRK2 mutation. (A) Ciliogenesis was measured in R1441C-LRRK2-KI MEFs that were serum-starved for 24 h -/+ the indicated peptides followed by immunocytochemistry for γ-tubulin (ciliary base, Alexa647-labeled secondary antibody, colored red), Arl13b (ciliary marker, Alexa594-labeled secondary antibody, colored pseudo-green) and DAPI (blue) (scale bar, 20 μm). The RIP peptides tested rescued ciliogenesis defects in a dose-dependent manner and almost to the same extent as the kinase inhibitor MLi-2 (200 nM, 24 h). (B) Quantification of ciliogenesis scored from 200 cells per condition and experiment (n=2 independent experiments each) demonstrates a dose-dependent effect on restoration of ciliogenesis defects.

### 3.5 Peptides Reduce Centrosomal Cohesion Defects in Cells

Since increased phosphorylation of Rab8 and Rab10 by LRRK2 is also implicated in centrosomal cohesion defects, we wanted to determine whether these effects could be restored with RIP peptides (**Figure 5A, B**). To test this, 1-10 μM of individual RIP peptides were tested using R1441C-knock-in MEF cells. Cells were treated with RIP peptides for 4 hours in medium containing serum prior to imaging using immunocytochemistry to visualize γ-tubulin (centrosomal marker) and DAPI. For each condition and experiment, cohesion was assessed using 100 cells with two centrosomes [27]. From this experiment, it was found that all RIP peptides tested could restore centrosomal cohesion in a dose-dependent manner and to a level comparable to the LRRK2 inhibitor, MLi-2. Thus, several RIP peptides were identified that could restore both ciliogenesis defects and centrosomal cohesion defects in the presence of a disease-relevant mutant form of LRRK2.

**Figure 5.**
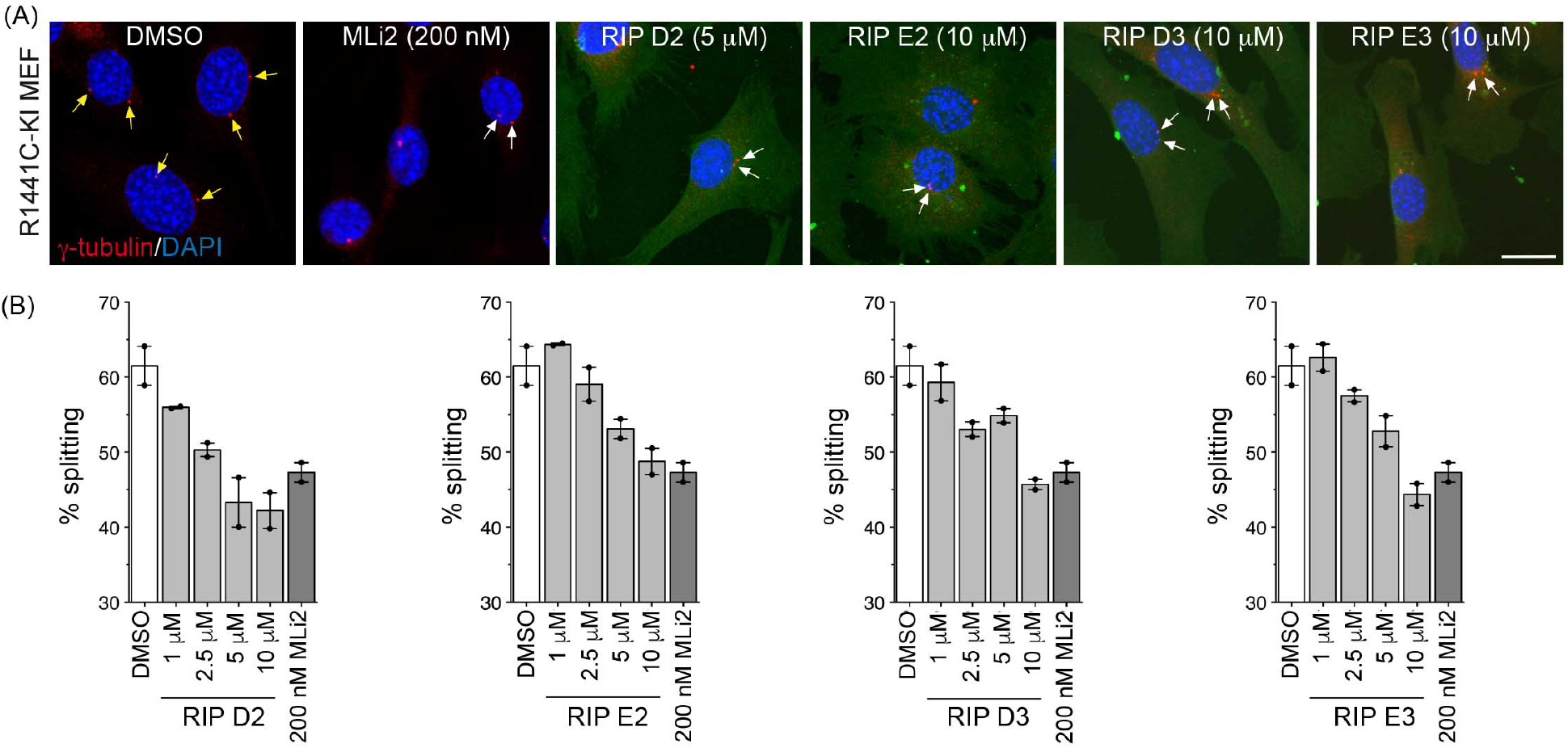
Constrained peptides rescue centrosomal splitting phenotype in a cell line harboring a PD-associated LRRK2 mutation. (A) R1441C-LRRK2-KI-MEFs were treated with RIP peptides in medium with serum for 4 h prior to imaging. Immunocytochemistry staining was performed for γ-tubulin (red), DAPI (blue), and FAM-labeled peptide fluorescence (green) (scale bar, 20 μm). Yellow arrows point to two split centrosomes, and white arrows to two attached centrosomes. (B) Quantification of centrosomal splitting for RIP peptides. For each condition and experiment, cohesion was assessed from 100 cells with two centrosomes (n=2 independent experiments each). All peptides tested were found to rescue centrosomal cohesion defects in a dose-dependent manner.

## CONCLUSION

Parkinson’s Disease (PD) is the second most commonly occurring neurodegenerative disorder worldwide and remains uncurable [30]. Mutations in LRRK2 are the most common cause of familial PD, resulting in enhanced LRRK2 kinase activity [1]. Although multiple treatments are in clinical use to control the motor symptoms of PD, none are yet approved that specifically target LRRK2 and its causative effects on disease progression. Tremendous efforts have been put forth toward developing LRRK2 kinase inhibitors and clinical trials are currently underway for some LRRK2-targeting agents [31]. However, numerous concerns including lung aberrations have been noted for several LRRK2 kinase inhibitors, highlighting the need to explore alternative strategies for LRRK2 inhibition.

In this study, we developed constrained peptides that were designed to inhibit a PPI that is known to be upregulated by aberrant LRRK2 activity: RILPL2 and phosphorylated Rab8a. Several of these RIP peptides (RIP D2, RIP E2, RIP D3, and RIP E3) were found to bind their intended RILPL2 target and could downregulate pathogenic effects of LRRK2, namely ciliogenesis and centrosomal cohesion defects. Further, these effects were comparable to directly inhibiting the kinase activity of LRRK2 itself via inhibition by MLi-2. This demonstrates that a non-catalytic approach to targeting specific pathways upregulated by LRRK2 may serve as an alternative strategy to downregulate various consequences of increased LRRK2 activity. Further, this general strategy may serve to circumvent the toxicities induced by many LRRK2 kinase inhibitors. To our knowledge, there are currently no other inhibitors that specifically target downstream signaling effects of LRRK2, and thus these constrained peptides may serve as powerful tool compounds for dissecting the effects of LRRK2 in PD.

There are still many unknowns regarding the role of phosphorylated Rab proteins and their interactions with phospho-specific effector proteins in the context of elevated LRRK2 kinase activity. While the phosphomimetic RIP peptides in this study were derived from Rab8a and did not appear to interact with JIP4, both JIP3 and JIP4 have been shown to interact with phosphorylated Rab10 [26]. Thus, an analogous peptide derived from Rab10 may have very different biological actions as compared to the RIP peptides described in this work. This may provide an interesting chemical biology strategy to dissect the unique cellular roles of different phosphorylated Rabs downstream of LRRK2. Further, there is considerable sequence diversity in the switch II region amongst the Rab superfamily, and this will dictate target specificity and likely influence target affinity. Further development of Rab-specific mimics may serve as useful tools to help unravel the roles of individual phospho-Rab proteins within a cellular context.

## ACKNOWLEDGEMENTS

This work was supported by the Michael J. Fox Foundation for Parkinson’s Research (MJFF-8068.04 for EJK and MJFF-019358 for SH).

## REFERENCES

[1] J.H. Kluss, A. Mamais, M.R. Cookson, LRRK2 links genetic and sporadic Parkinson’s disease, Biochem Soc Trans, 47 (2019) 651–661.

[2] C. Schulte, T. Gasser, Genetic basis of Parkinson’s disease: inheritance, penetrance, and expression, Appl Clin Genet, 4 (2011) 67–80.

[3] I.N. Rudenko, M.R. Cookson, Heterogeneity of leucine-rich repeat kinase 2 mutations: genetics, mechanisms and therapeutic implications, Neurotherapeutics, 11 (2014) 738–750.

[4] M. Steger, F. Tonelli, G. Ito, P. Davies, M. Trost, M. Vetter, S. Wachter, E. Lorentzen, G. Duddy, S. Wilson, M.A. Baptista, B.K. Fiske, M.J. Fell, J.A. Morrow, A.D. Reith, D.R. Alessi, M. Mann, Phosphoproteomics reveals that Parkinson’s disease kinase LRRK2 regulates a subset of Rab GTPases, Elife, 5 (2016) e12813.

[5] Z. Liu, N. Bryant, R. Kumaran, A. Beilina, A. Abeliovich, M.R. Cookson, A.B. West, LRRK2 phosphorylates membrane-bound Rabs and is activated by GTP-bound Rab7L1 to promote recruitment to the trans-Golgi network, Hum Mol Genet, 27 (2018) 385–395.

[6] X. Deng, N. Dzamko, A. Prescott, P. Davies, Q. Liu, Q. Yang, J.D. Lee, M.P. Patricelli, T.K. Nomanbhoy, D.R. Alessi, N.S. Gray, Characterization of a selective inhibitor of the Parkinson’s disease kinase LRRK2, Nat Chem Biol, 7 (2011) 203–205.

[7] J. Zhang, X. Deng, H.G. Choi, D.R. Alessi, N.S. Gray, Characterization of TAE684 as a potent LRRK2 kinase inhibitor, Bioorg Med Chem Lett, 22 (2012) 1864–1869.

[8] A.D. Reith, P. Bamborough, K. Jandu, D. Andreotti, L. Mensah, P. Dossang, H.G. Choi, X. Deng, J. Zhang, D.R. Alessi, N.S. Gray, GSK2578215A; a potent and highly selective 2-arylmethyloxy-5-substitutent-N-arylbenzamide LRRK2 kinase inhibitor, Bioorg Med Chem Lett, 22 (2012) 5625–5629.

[9] N. Ramsden, J. Perrin, Z. Ren, B.D. Lee, N. Zinn, V.L. Dawson, D. Tam, M. Bova, M. Lang, G. Drewes, M. Bantscheff, F. Bard, T.M. Dawson, C. Hopf, Chemoproteomics-based design of potent LRRK2-selective lead compounds that attenuate Parkinson’s disease-related toxicity in human neurons, ACS Chem Biol, 6 (2011) 1021–1028.

[10] H.G. Choi, J. Zhang, X. Deng, J.M. Hatcher, M.P. Patricelli, Z. Zhao, D.R. Alessi, N.S. Gray, Brain Penetrant LRRK2 Inhibitor, ACS Med Chem Lett, 3 (2012) 658–662.

[11] M.A. Baptista, K.D. Dave, M.A. Frasier, T.B. Sherer, M. Greeley, M.J. Beck, J.S. Varsho, G.A. Parker, C. Moore, M.J. Churchill, C.K. Meshul, B.K. Fiske, Loss of leucine-rich repeat kinase 2 (LRRK2) in rats leads to progressive abnormal phenotypes in peripheral organs, PLoS One, 8 (2013) e80705.

[12] R.N. Fuji, M. Flagella, M. Baca, M.A. Baptista, J. Brodbeck, B.K. Chan, B.K. Fiske, L. Honigberg, A.M. Jubb, P. Katavolos, D.W. Lee, S.C. Lewin-Koh, T. Lin, X. Liu, S. Liu, J.P. Lyssikatos, J. O’Mahony, M. Reichelt, M. Roose-Girma, Z. Sheng, T. Sherer, A. Smith, M. Solon, Z.K. Sweeney, J. Tarrant, A. Urkowitz, S. Warming, M. Yaylaoglu, S. Zhang, H. Zhu, A.A. Estrada, R.J. Watts, Effect of selective LRRK2 kinase inhibition on nonhuman primate lung, Sci Transl Med, 7 (2015) 273ra215.

[13] M. Steger, F. Diez, H.S. Dhekne, P. Lis, R.S. Nirujogi, O. Karayel, F. Tonelli, T.N. Martinez, E. Lorentzen, S.R. Pfeffer, D.R. Alessi, M. Mann, Systematic proteomic analysis of LRRK2-mediated Rab GTPase phosphorylation establishes a connection to ciliogenesis, Elife, 6 (2017) e31012.

[14] A.H. Hutagalung, P.J. Novick, Role of Rab GTPases in membrane traffic and cell physiology, Physiol Rev, 91 (2011) 119–149.

[15] N.A. Guadagno, C. Progida, Rab GTPases: Switching to Human Diseases, Cells, 8 (2019).

[16] H. Stenmark, Rab GTPases as coordinators of vesicle traffic, Nat Rev Mol Cell Biol, 10 (2009) 513–525.

[17] H.S. Dhekne, I. Yanatori, R.C. Gomez, F. Tonelli, F. Diez, B. Schule, M. Steger, D.R. Alessi, S.R. Pfeffer, A pathway for Parkinson’s Disease LRRK2 kinase to block primary cilia and Sonic hedgehog signaling in the brain, Elife, 7 (2018) e40202.

[18] J.R. Schaub, T. Stearns, The Rilp-like proteins Rilpl1 and Rilpl2 regulate ciliary membrane content, Mol Biol Cell, 24 (2013) 453–464.

[19] A.J. Lara Ordonez, B. Fernandez, E. Fdez, M. Romo-Lozano, J. Madero-Perez, E. Lobbestael, V. Baekelandt, A. Aiastui, A. Lopez de Munain, H.L. Melrose, L. Civiero, S. Hilfiker, RAB8, RAB10 and RILPL1 contribute to both LRRK2 kinase-mediated centrosomal cohesion and ciliogenesis deficits, Hum Mol Genet, 28 (2019) 3552–3568.

[20] J. Madero-Perez, E. Fdez, B. Fernandez, A.J. Lara Ordonez, M. Blanca Ramirez, P. Gomez-Suaga, D. Waschbusch, E. Lobbestael, V. Baekelandt, A.C. Nairn, J. Ruiz-Martinez, A. Aiastui, Lopez de Munain, P. Lis, T. Comptdaer, J.M. Taymans, M.C. Chartier-Harlin, A. Beilina, A. Gonnelli, M.R. Cookson, E. Greggio, S. Hilfiker, Parkinson disease-associated mutations in LRRK2 cause centrosomal defects via Rab8a phosphorylation, Mol Neurodegener, 13 (2018) 3.

[21] A.J. Lara Ordonez, R. Fasiczka, B. Fernandez, Y. Naaldijk, E. Fdez, M. Blanca Ramirez, S. Phan, D. Boassa, S. Hilfiker, The LRRK2 signaling network converges on a centriolar phospho-Rab10/RILPL1 complex to cause deficits in centrosome cohesion and cell polarization, Biol Open, 11 (2022) bio059468.

[22] H.S. Dhekne, I. Yanatori, E.G. Vides, Y. Sobu, F. Diez, F. Tonelli, S.R. Pfeffer, LRRK2-phosphorylated Rab10 sequesters Myosin Va with RILPL2 during ciliogenesis blockade, Life Sci Alliance, 4 (2021).

[23] B. Fernandez, A.J. Lara Ordonez, E. Fdez, E. Mutez, T. Comptdaer, C. Leghay, A. Kreisler, Simonin, L. Vandewynckel, L. Defebvre, A. Destee, S. Bleuse, J.M. Taymans, M.C. Chartier-Harlin, S. Hilfiker, Centrosomal cohesion deficits as cellular biomarker in lymphoblastoid cell lines from LRRK2 Parkinson’s disease patients, Biochem J, 476 (2019) 2797–2813.

[24] Y. Naaldijk, B. Fernández, R. Fasiczka, E. Fdez, C. Leghay, I. Croitoru, J.B. Kwok, Y. Boulesnane, A. Vizeneux, E. Mutez, C. Calvez, A. Destée, J.-M. Taymans, A.V. Aragon, A.B. Yarza, S. Padmanabhan, M. Delgado, R.N. Alcalay, Z. Chatterton, N. Dzamko, G. Halliday, J. Ruiz-Martìnez, M.-C. Chartier-Harlin, S. Hilfiker, A potential patient stratification biomarker for Parkinso’s disease based on LRRK2 kinase-mediated centrosomal alterations in peripheral blood-derived cells, bioRxiv, (2023) 2023.2004.2011.536367.

[25] S.S. Khan, Y. Sobu, H.S. Dhekne, F. Tonelli, K. Berndsen, D.R. Alessi, S.R. Pfeffer, Pathogenic LRRK2 control of primary cilia and Hedgehog signaling in neurons and astrocytes of mouse brain, Elife, 10 (2021) e67900.

[26] D. Waschbusch, E. Purlyte, P. Pal, E. McGrath, D.R. Alessi, A.R. Khan, Structural Basis for Rab8a Recruitment of RILPL2 via LRRK2 Phosphorylation of Switch 2, Structure, 28 (2020) 406–417 e406.

[27] E. Fdez, R. Fasiczka, A.J. Lara Ordonez, B. Fernandez, Y. Naaldijk, S. Hilfiker, Protocol to measure centrosome cohesion deficits mediated by pathogenic LRRK2 in cultured cells using confocal microscopy, STAR Protoc, 4 (2023) 102024.

[28] Y. Sobu, P.S. Wawro, H.S. Dhekne, W.M. Yeshaw, S.R. Pfeffer, Pathogenic LRRK2 regulates ciliation probability upstream of tau tubulin kinase 2 via Rab10 and RILPL1 proteins, Proc Natl Acad Sci U S A, 118 (2021).

[29] E. Fdez, J. Madero-Perez, A.J. Lara Ordonez, Y. Naaldijk, R. Fasiczka, A. Aiastui, J. Ruiz-Martinez, A. Lopez de Munain, S.A. Cowley, R. Wade-Martins, S. Hilfiker, Pathogenic LRRK2 regulates centrosome cohesion via Rab10/RILPL1-mediated CDK5RAP2 displacement, iScience, 25 (2022) 104476.

[30] S. Bandres-Ciga, M. Diez-Fairen, J.J. Kim, A.B. Singleton, Genetics of Parkinson’s disease: An introspection of its journey towards precision medicine, Neurobiol Dis, 137 (2020) 104782.

[31] S. Azeggagh, D.C. Berwick, The development of inhibitors of leucine-rich repeat kinase 2 (LRRK2) as a therapeutic strategy for Parkinson’s disease: the current state of play, Br J Pharmacol, 179 (2022) 1478–1495.

