## Supplementary Information for "Targeting Rab-RILPL Interactions as a Strategy to Downregulate Pathogenic LRRK2 in Parkinson’s Disease"

Supporting Figures

Figure S1. ESI-MS Analysis of Native Peptide; FAM and Biotin Labels
Figure S2. ESI-MS Analysis of RIP D2; FAM and Biotin Labels
Figure S3. ESI-MS Analysis of RIP E2; FAM and Biotin Labels
Figure S4. ESI-MS Analysis of RIP D3; FAM and Biotin Labels
Figure S5. ESI-MS Analysis of RIP E3; FAM and Biotin Labels
Figure S6. Large Field View of Cell Uptake
Figure S7. Proteolytic Stability

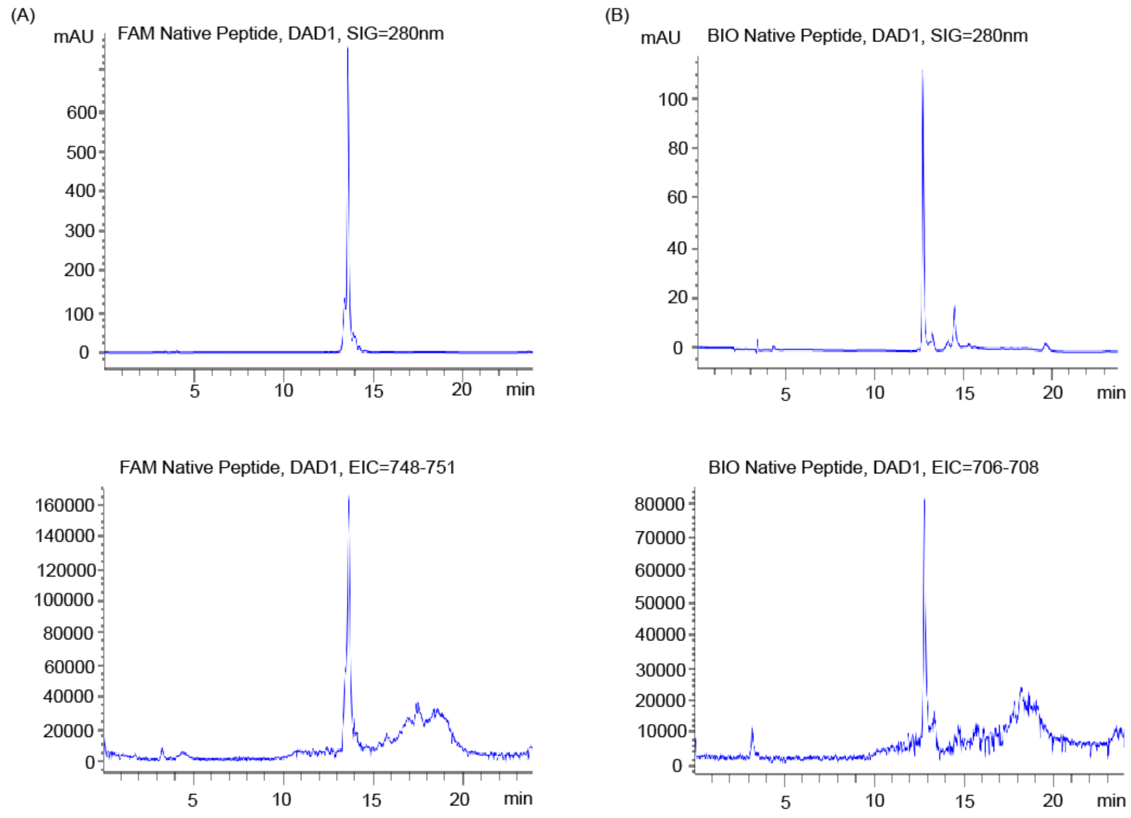


**Figure  S1. ESI-MS Analysis of Native Peptide; FAM and Biotin Labels.** (A) FAM-labeled Native Peptide ESI-MS; molecular weight: 2250.0 g/mol (Expected:2250.6 g/mol) and (B) Biotin-Labeled Native Peptide ESI-MS; molecular weight: 2118.0 g/mol (Expected:2118.5 g/mol)

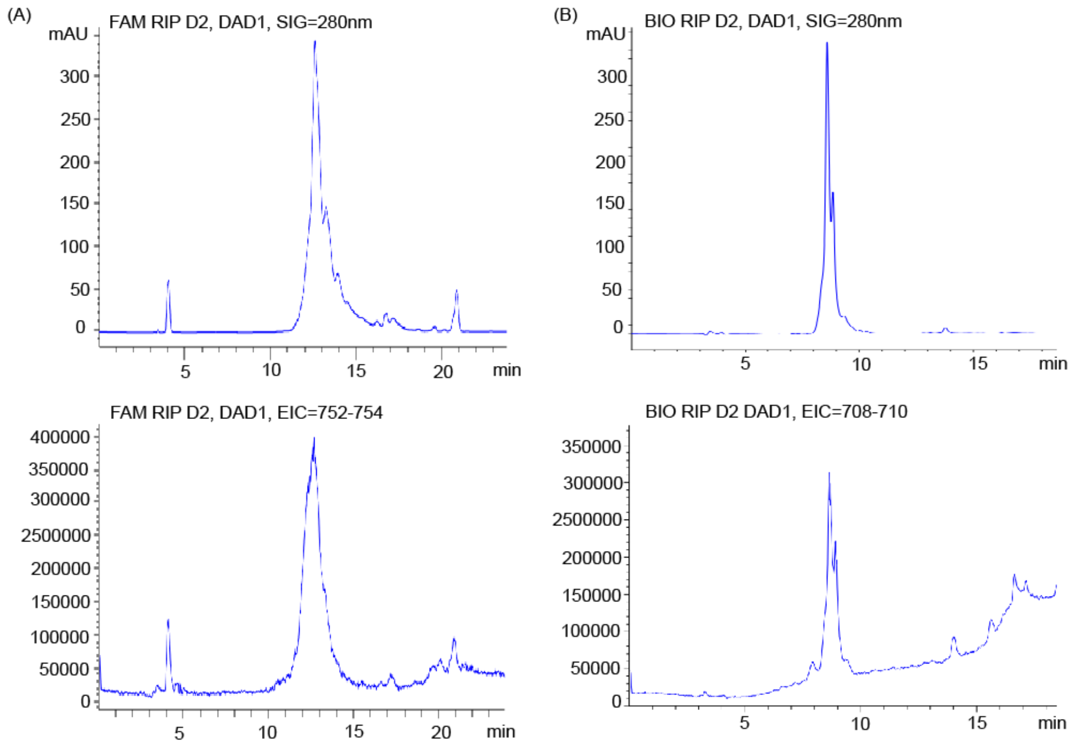


**Figure S2. ESI-MS Analysis of RIP D2; FAM Label.** (A) FAM-labeled RIP D2 Peptide ESI-MS; molecular weight: 2256.9 g/mol (Expected:2257.6 g/mol) and (B) Biotin-Labeled RIP D2 Peptide ESI-MS; molecular weight: 2124.0 g/mol (Expected:2124.2 g/mol)

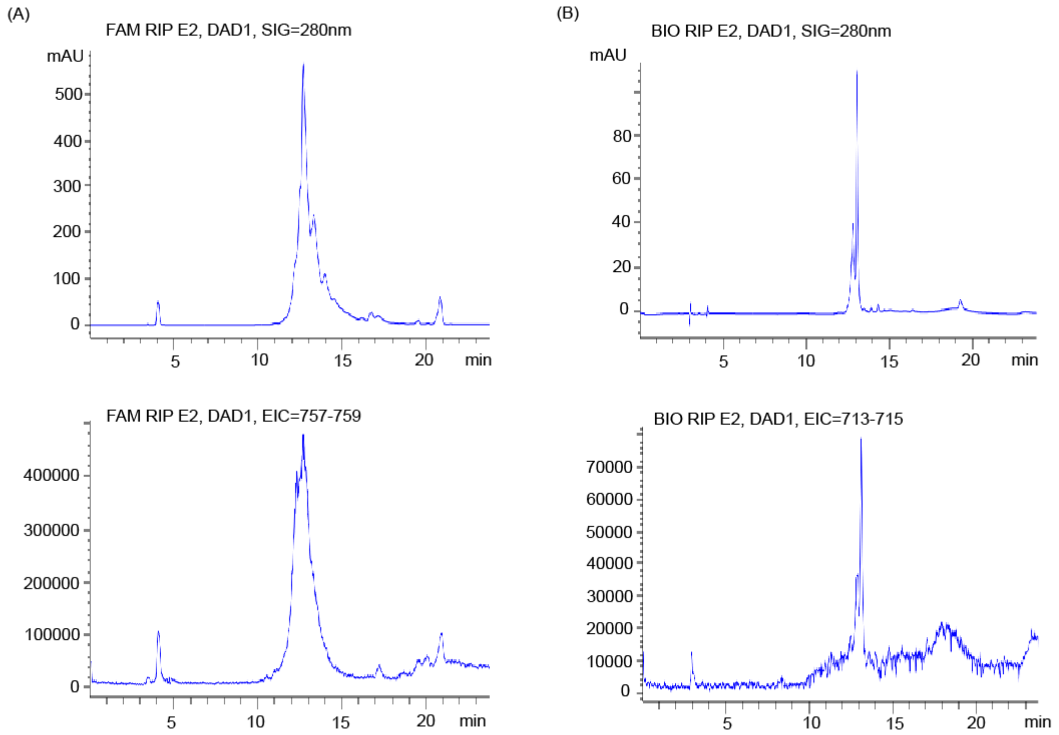


**Figure S3. ESI-MS Analysis of RIP E2; FAM and Biotin Labels.** (A) FAM-labeled RIP E2 Peptide ESI-MS; molecular weight: 2270.7 g/mol (Expected:2271.6) and (B) Biotin-Labeled RIP E2 Peptide ESI-MS; molecular weight: 2139.0 g/mol (Expected:2139.6 g/mol)

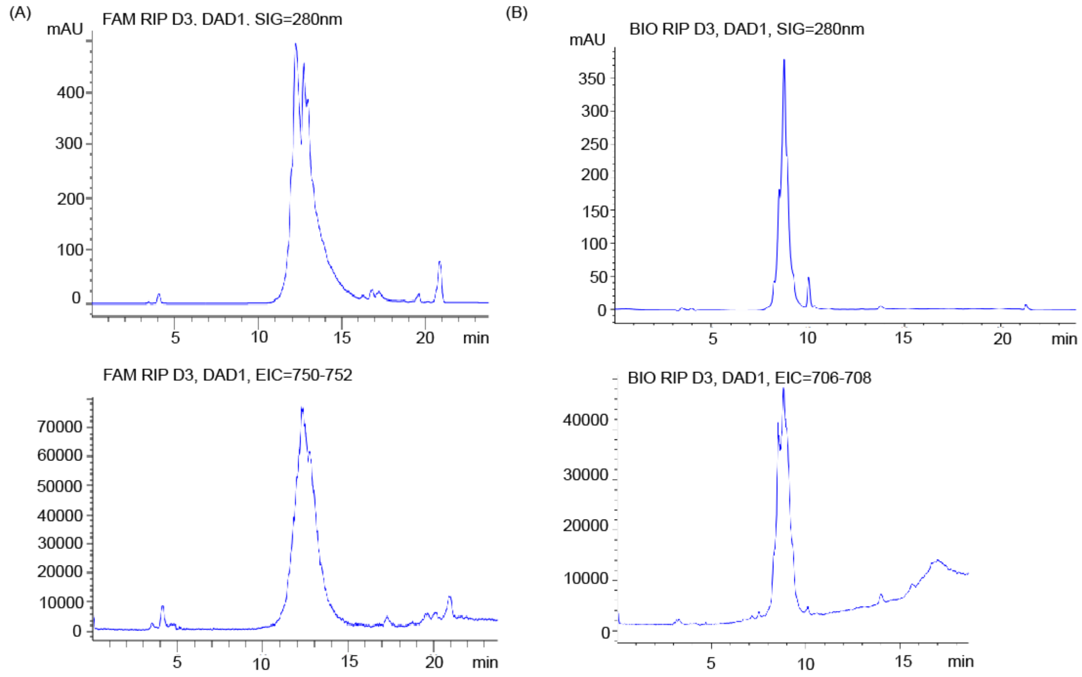


**Figure S4. ESI-MS Analysis of RIP D3; FAM and Biotin Labels.** (A) FAM-labeled RIP D3 Peptide ESI-MS; molecular weight: 2250.0 g/mol (Expected:2250.6 g/mol) and (B) Biotin-Labeled RIP D3 Peptide ESI-MS; molecular weight: 2118.0 g/mol (Expected:2118.6 g/mol)

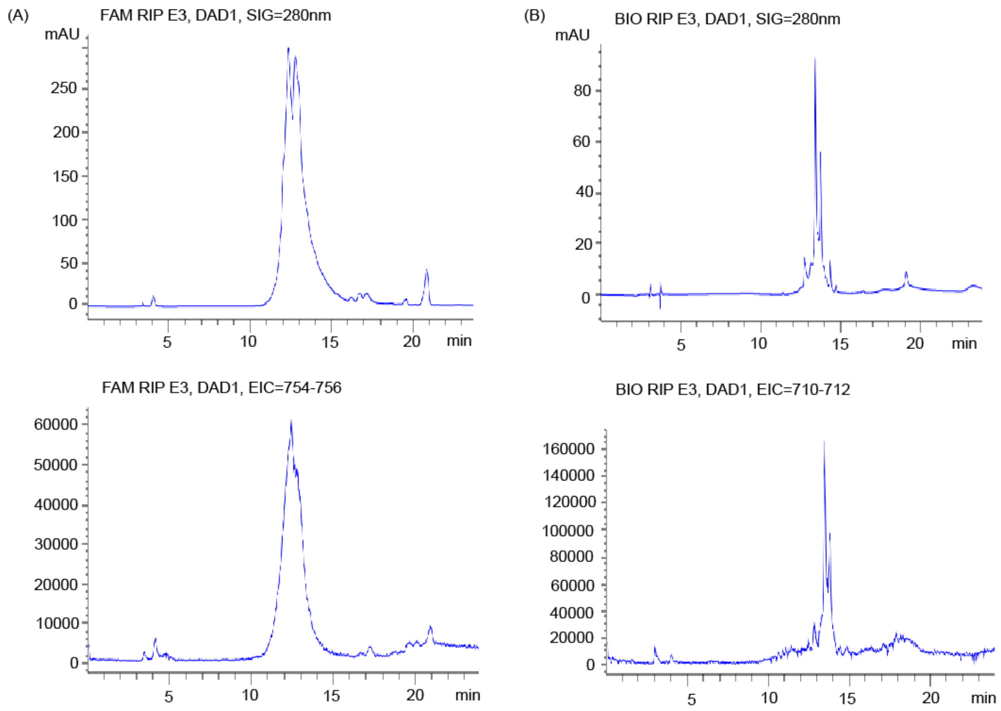


**Figure S5. ESI-MS Analysis of RIP E3; FAM and Biotin Labels.** (A) FAM-labeled RIP E3 Peptide ESI-MS; molecular weight: 2264.1 g/mol (Expected:2264.6 g/mol) and (B) Biotin-Labeled RIP E3 Peptide ESI-MS; molecular weight: 2132.1 g/mol (Expected:2132.5 g/mol)

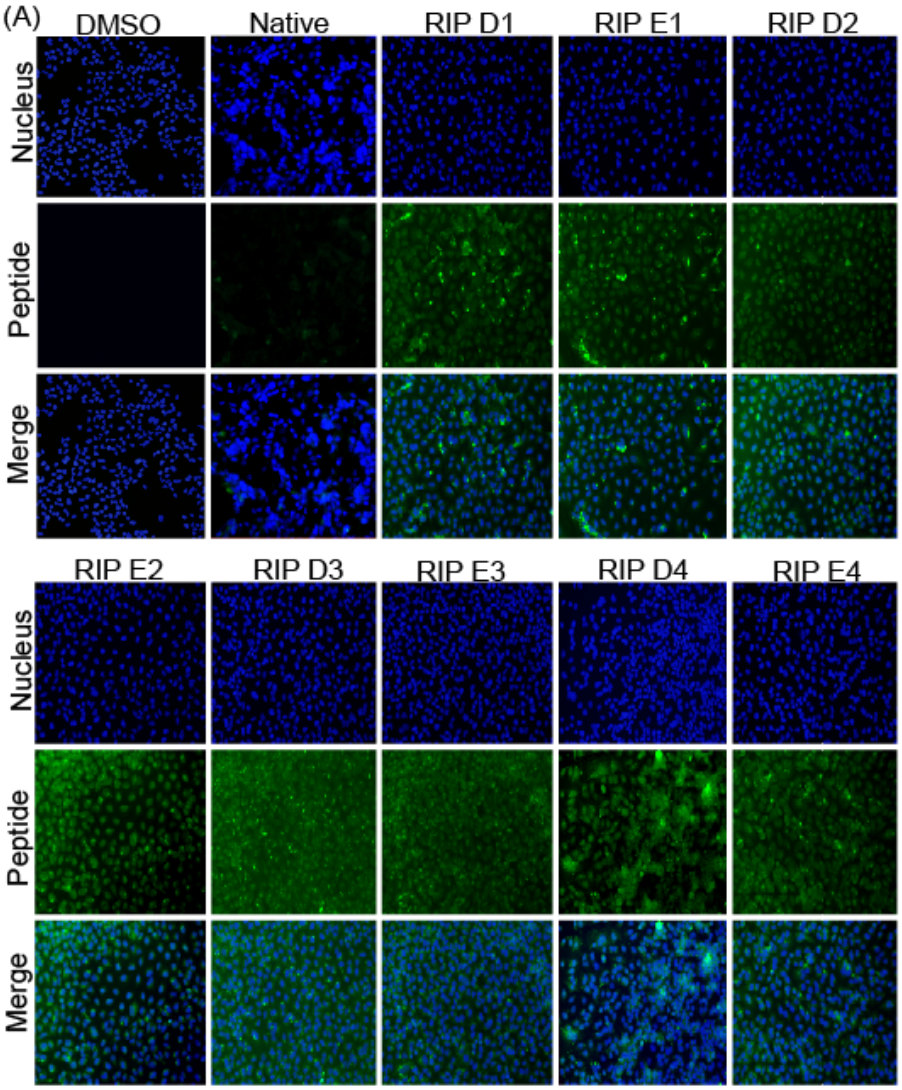


**Figure S6. Large Field View of Cell Uptake.** (A) Large field view of FAM-labeled peptide uptakes. A549 cells were treated with 2.5 μM of each FAM-labeled peptide for 8 hours prior to fluorescent imaging. Over 200 cells were studied and captured per peptide treatment. The unstapled native peptide sequence did not permeate cells, while the stapled peptides permeated cells to varying degrees.

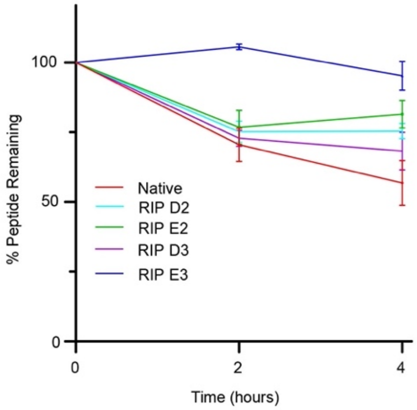


**Figure S7. Proteolytic Stability.** (A) The native peptide, RIP D2, RIP E2, RIP D3, and RIP E3 were individually incubated in a mouse serum cocktail at 37 °C for 4 hours, with mass spectrometry measurements taken every 2 hours. While the native unstapled peptide was degraded by roughly 50% over this time course, the RIP peptides maintained between 75%-95% integrity over the same time frame. The graph represents three independent experiments with error bars indicating standard deviation.
